# Elovl2 but not Elovl5 is essential for the biosynthesis of docosahexaenoic acid (DHA) in zebrafish: insight from a comparative gene knockout study

**DOI:** 10.1101/2020.06.03.132951

**Authors:** Chengjie Liu, Ding Ye, Houpeng Wang, Mudan He, Yonghua Sun

## Abstract

Teleost fish can synthesize one of the major omega-3 long-chain polyunsaturated fatty acids (n-3 LC-PUFAs), docosahexaenoic acid (DHA, 22:6n-3), from dietary α-linolenic acid (ALA; 18:3n-3), via elongase of very long chain fatty acid (Elovl) and fatty acid desaturase (Fads). However, it remains unclear which elongase is responsible for the endogenous synthesis of DHA. Here in this study, the knockout models of the two major elongases, Elovl2 and Elovl5, were generated by CRISPR/Cas9 approach in zebrafish and comparatively analyzed. The homozygous mutants were validated by Sanger sequencing, mutation-mediated PCR and whole-mount in situ hybridization analysis of the endogenous target genes. Compared with wildtype (WT) counterparts, the content of DHA was significantly reduced by 67.1% (p<0.05) in the adult liver and by 91.7% (p<0.01) in the embryo at 3 day-post-fertilization (dpf) of the *elovl2* mutant, but not of the *elovl5* mutant. Further study revealed that *elovl2* and *fads2* was upregulated by 9.9-fold (p<0.01) and 9.7-fold (p<0.01) in the *elovl5* mutant, and *elovl5* and *fads2* was upregulated by 15.1-fold (p<0.01) and 21.5-fold (p<0.01) in the *elovl2* mutant. Our study indicates that although both Elovl2 and Elovl5 have the elongase activity toward C20, the upregulation of *elovl2* could completely replace the genetic depletion of *elovl5*, but upregulation of *elovl5* could not compensate the endogenous deficiency of *elovl2* in mediating DHA synthesis. In conclusion, the endogenous synthesis of DHA in is mediated by Elovl2 but not Elovl5 in teleost, and a DHA-deficient genetic model of zebrafish has been generated.

## Introduction

Long-chain polyunsaturated fatty acids (LC-PUFAs), which possess 20 or more carbon atoms and contain two or more double bonds in their carbon chains, e.g. docosahexaenoic acid (DHA, 22:6n-3), are essential nutrients for neural development and health (Heird and Lapillonne, 2005). During the biosynthesis of LC-PUFAs, fatty acid desaturase (Fads) and elongase of very long chain fatty acid (Elovl) are critical enzymes for desaturation and elongation in LC-PUFA synthesis from dietary essential fatty acids, α-linolenic acid (ALA, 18:3n-3) and linoleic acid (LA, 18:2n-6) (Monroig et al., 2013, Guillou et al., 2010). The enzyme activities of two major elongases of different species, Elovl2 and Elovl5, have been analyzed mostly in in vitro systems, such as yeast or cultured cells (Leonard et al., 2002). Functional characterization assays in the yeast system revealed that the zebrafish Elovl5 and Elovl2 both have the ability to elongate C18-C22 PUFA substrates (Agaba et al., 2004, Monroig et al., 2009). Although the substrate specificities of Elovl2 and Elovl5 show some species-specific diversity, both Elovl2 and Elovl5 have been shown to be able to elongate C20 LC-PUFA substrates (Monroig et al., 2013, Monroig et al., 2016, Monroig et al., 2012). Therefore, whether the Elovl2 or Elovl5 elongase is mainly responsible for the endogenous biosynthesis of DHA needs to be clarified with gene knockout models.

In this study, by generating *elovl2* and *elovl5* knockout zebrafish models, we revealed that Elovl2 dominantly mediates elongation from eicosapentaenoic acid (EPA, 20:5n-3) to DHA in teleost fish.

## Materials and Methods

### Zebrafish strain

Wildtype (WT) zebrafish of the AB strain were maintained and raised at the China Zebrafish Resource Center of the National Aquatic Biological Resource Center (CZRC/NABRC, http://zfish.cn, Wuhan, China) according to the Zebrafish Book (Westerfield, 2000). The adult fish were fed daily with Artemia, which contain trace amounts of DHA (the LC-PUFA profile is provided in Table 1). The experiments involving zebrafish followed the Zebrafish Usage Guidelines of CZRC and were performed under the approval of the Institutional Animal Care and Use Committee of the Institute of Hydrobiology, Chinese Academy of Sciences, under protocol IHB2015-006.

**Table 1.** LC-PUFAs composition^1^ of total fatty acids extracted from *Artemia*

| Fatty acid | Mol% of total fatty acid |
| --- | --- |
| 18:2n-6 | 6.04±0.05 |
| 18:3n-6 | 0.16±0.01 |
| 20:2n-6 | 0.19±0.01 |
| 20:3n-6 | 0.14±0.01 |
| 20:4n-6 | 2.07±0.08 |
| 22:4n-6 | 0.01±0.00 |
| 18:3n-3 | 9.60±0.02 |
| 20:5n-3 | 19.60±0.20 |
| 22:5n-3 | 0.04±0.01 |
| 22:6n-3 | 0.48±0.02 |

<sup>1</sup> Values are means ± SEM; n = 3 samples.

### Generation and identification of genetic knock-out fish

Genetic mutants of *elovl2* and *elovl5* were generated by a CRISPR/Cas9-mediated knockout approach as previously described (Chang et al., 2013, Ye et al., 2019). Briefly, the guide RNAs (gRNAs) GGTTACCGTCTTCAGTGTC<u>AGG</u> (PAM sequence underlined), targeting the fourth exon of *elovl2*, and GGAGAAGTAATACCACCAC<u>AGG</u> (PAM sequence underlined) targeting the fifth exon of *elovl5*, were designed using CRISPRscan (http://www.crisprscan.org/) (Moreno-Mateos et al., 2015). The gRNAs were synthesized using gRNA-pMD19-T (CZRC Plasmid #CZP3) as the PCR template according to the reported method (Chang et al., 2013). The primers used for gRNAs synthesis showed in Table 2. The zebrafish codon-optimized *cas9* mRNA was transcribed in vitro from pCS2-nzcas9n (CZRC Plasmid #CZP13) (Jao et al., 2013). gRNA (150 pg/embryo) and cas9 mRNA (500 pg/embryo) were co-injected into zebrafish embryos at 1-cell stage as previously described (Zhang et al., 2020). The mutations were identified by Sanger sequencing of the PCR products covering target sites using the primer pairs *elovl2*-testF/R and *elovl5*-testF/R (Table 2). For screening of the *elovl2* and *elovl5* homozygous mutant, genomic DNA extracted from zebrafish tail fin was used as a template for PCR reactions using the WT allele-specific and mutant allele-specific primer pairs in Table 2 (e2F1/e2R and e2F2/e2R for *elovl2*, e5F1/e5R and e5F2/e5R for *elovl5*), followed by agarose gel electrophoresis of the PCR products to identify the different genotypes.

**Table 2.**
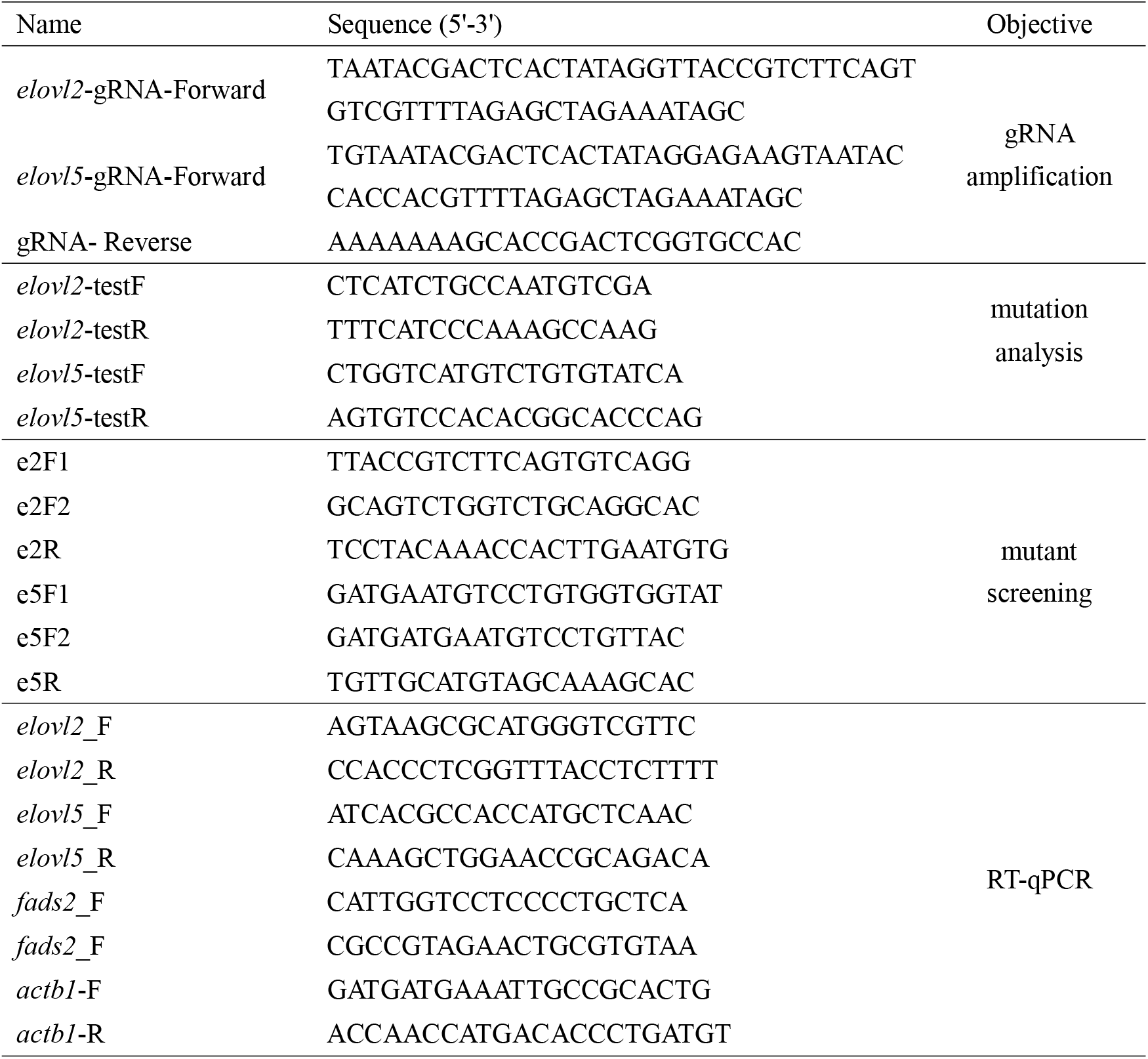
List of primers used in present study

### LC-PUFA analysis

The livers of adult zebrafish at 3 months post-fertilization (mpf), zebrafish embryos at 3 days post-fertilization (dpf) and fish diet were used for lipid extraction. The quantification of fatty acids was performed by gas chromatography-mass spectrometry (GC-MS) according to our previous studies (Zhang et al., 2019, Pang et al., 2014). LC-PUFAs were analyzed in three independent samples, with each sample containing liver tissues of three fishes, 100 embryos or 0.1~0.2 g fish diet of brine shrimp.

### Whole-mount *in situ* hybridization

Whole-mount *in situ* hybridization (WISH) of embryos at 3 dpf was performed as previously described (Ye et al., 2019). The signal was developed by NBT-BCIP kit. Before imaging, the embryos were soaked in the 100% glycerol overnight at 4 °C. The images were acquired using a Leica Z16 APO macroscope with a Leica DFC 450FX CCD.

### Reverse-transcription PCR and reverse-transcription quantitative PCR

RNA was extracted from livers of 3 mpf zebrafish by Trizol (Invitrogen) and the cDNA was synthesized by RevertAid cDNA Synthesis Kit (Thermo). For both reverse-transcription PCR (RT-PCR) and reverse-transcription quantitative PCR (RT-qPCR) analysis of *elovl2* and *elovl5*, cDNA was amplified with the primers *elovl2*_F/ *elovl2*_R for *elovl2* (product size: 135 bp) and *elovl5*_F/ *elovl5*_R for *elovl5* (product size: 140 bp), and *actb1* was used as the internal control. The primers were listed in Table 2. For RT-qPCR analysis, real-time PCR was performed in a BioRad CFX Connect Real-Time System using SYBR Green mix (BioRad) according to the MIQE (Minimum Information for Publication of Quantitative Real-Time PCR Experiments) guidelines (Taylor et al., 2010). The expression levels of mRNA were calculated based on the −ΔΔCT method according to a previous study (Pfaffl, 2001). Each RT-qPCR analysis was repeated in triplicates. The primers sequences for RT-qPCR were listed in Table 2.

### Statistical analysis

The results are expressed as the mean ± SEM. Differences between 2 groups were tested by Student’s t-test. Differences were considered significant at *P* < 0.05.

## Results

### Generation of zebrafish *elovl2* and *elovl5* mutants by CRISPR/Cas9

The dynamic expression of *elovl2* and *elovl5* during zebrafish embryogenesis has been carefully studied previously (Monroig et al., 2009). Here, we found that both genes were highly expressed in liver and intestine, and they showed moderate expression levels ovary and testis of the adults (Fig. 1a). Moreover, they both showed weak expression in the brain and gills, with *elovl2* showing higher expression than *elovl5* in the brain, whereas *elovl5* showed higher expression than *elovl2* in the gills (Fig. 1a). By WISH analysis, both *elovl2* and *elovl5* are specifically expressed in the liver primordium in the zebrafish embryos at 3 dpf (Fig. 1b). These data suggested that the liver and intestine might be the main organs for LC-PUFA synthesis in zebrafish.

**Fig. 1.**
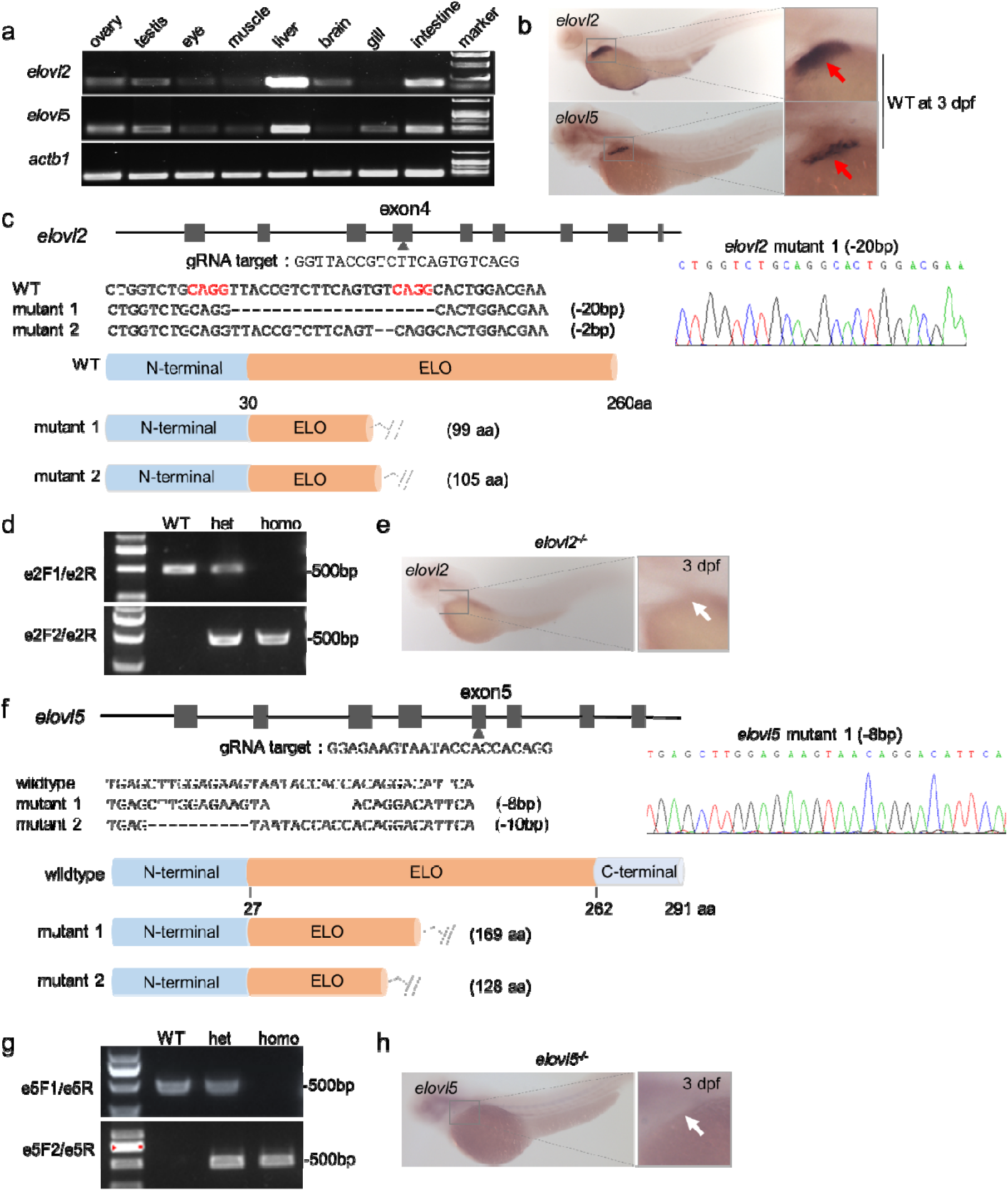
Generation and validation of *elovl2* and *elovl5* mutant zebrafish. (a) RT-PCR analysis of *elovl2* and *elovl5* in different tissues. (b) Expression patterns of *elovl5* and *elovl2* in WT embryos (3 dpf) examined by WISH. The red arrows showed the signals of *elovl2* and *elovl5* in the liver primordium region. (c) Diagram showing the gRNA target of *elovl2*, the genotypes and the length and structure of the predicted mutant proteins. Sanger sequencing was used to detect the *elovl2* mutation type 1 with 20 bp deletion. (d) Identification of WT, *elovl2* heterozygous (het), and *elovl2* homozygous (homo) fish with PCR primer pairs, e2F1/e2R and e2F2/e2R. (e) Expression pattern of *elovl2* in *elovl2*^*−/−*^ embryos (3 dpf) examined by WISH. The white arrow showed that the signal of *elovl2* was not detected in the liver primordium. (f) Diagram showing the gRNA target of *elovl5*, the genotypes and the length and structure of the predicted mutant proteins. Sanger sequencing was used to detect the *elovl5* mutation type 1 with 8 bp deletion. (g) Identification of WT, *elovl5* heterozygous (het), and *elovl5* homozygous (homo) fish with PCR primer pairs, e5F1/e5R and e5F2/e5R. (h) Expression pattern of *elovl5* in the *elovl5*^*−/−*^ embryos (3 dpf) examined by WISH. The white arrow showed that the signal of *elovl5* was not detected in the liver primordium.

To verify the in vivo function of these two elongases in fish, we then generated *elovl2* and *elovl5* mutants via the CRISPR/Cas9 approach. Two alleles were generated for the *elovl2* mutants, with 20 bp deletion or 2 bp deletion in the coding sequence of the elongase domain (Fig. 1c). Sanger sequencing showed that the 20 bp sequence was deleted in the *elovl2* homozygous mutant (Fig. 1c). Therefore, we designed WT-specific (e2F1/e2R) and mutant-specific (e2F2/e2R) primer pairs to screen the WT, heterozygote and homozygote of *elovl2* mutants (Fig. 1d). In the *elovl2*^*−/−*^ embryo at 3 dpf, WISH analysis showed that the transcription of endogenous *elovl2* totally disappeared (Fig. 1e), further indicating the genetic depletion of *elovl2*.

Similarly, two alleles were generated for the *elovl5* mutants, with 8 bp deletion or 10 bp deletion in the coding sequence of the elongase domain (Fig. 1f). Sanger sequencing confirmed that the 8 bp sequence was deleted in the *elovl5* homozygous mutant (Fig. 1f). We also designed WT-specific (e5F1/e5R) and mutant-specific (e5F2/e5R) primer pairs to screen the WT, heterozygote and homozygote of *elovl5* mutants (Fig. 1g). WISH analysis showed that the transcription of endogenous *elovl5* totally disappeared in the *elovl5*^*−/−*^ embryo at 3 dpf (Fig. 1h), further indicating the genetic depletion of *elovl5*. All these results suggest that both genes were effectively mutated in the mutant fish and that their transcribed mRNAs would be degraded due to nonsense mRNA decay mechanism (Popp and Maquat, 2016).

### PUFA analysis and gene expression analysis of *evlol2* and *elovl5* mutants

Then we compared the fatty acids composition in the livers of WT and two mutant zebrafish at adult stage. Interestingly, the amount of DHA in the liver of *elovl2*^*−/−*^ zebrafish (1.45±0.09%) was decreased by 67.1% (p<0.05), in comparison with that in the WT (4.42±0.93%). Whereas, the amount of EPA, the synthetic precursor of DHA and 22:5n-3, was increased by 48% (p<0.01) in the *elovl2*^*−/−*^ zebrafish (19.49±0.57%), compared with that in WT (13.15±0.77%) (Fig. 2a). In contrast, there was no significant difference in the contents of EPA and DHA between the WT and *elovl5*^*−/−*^ zebrafish (Fig. 2a). These results suggest that Elovl2 plays a dominant role in endogenous synthesis of DHA, while Elovl5 is not essential for DHA synthesis in zebrafish.

**Fig. 2.**
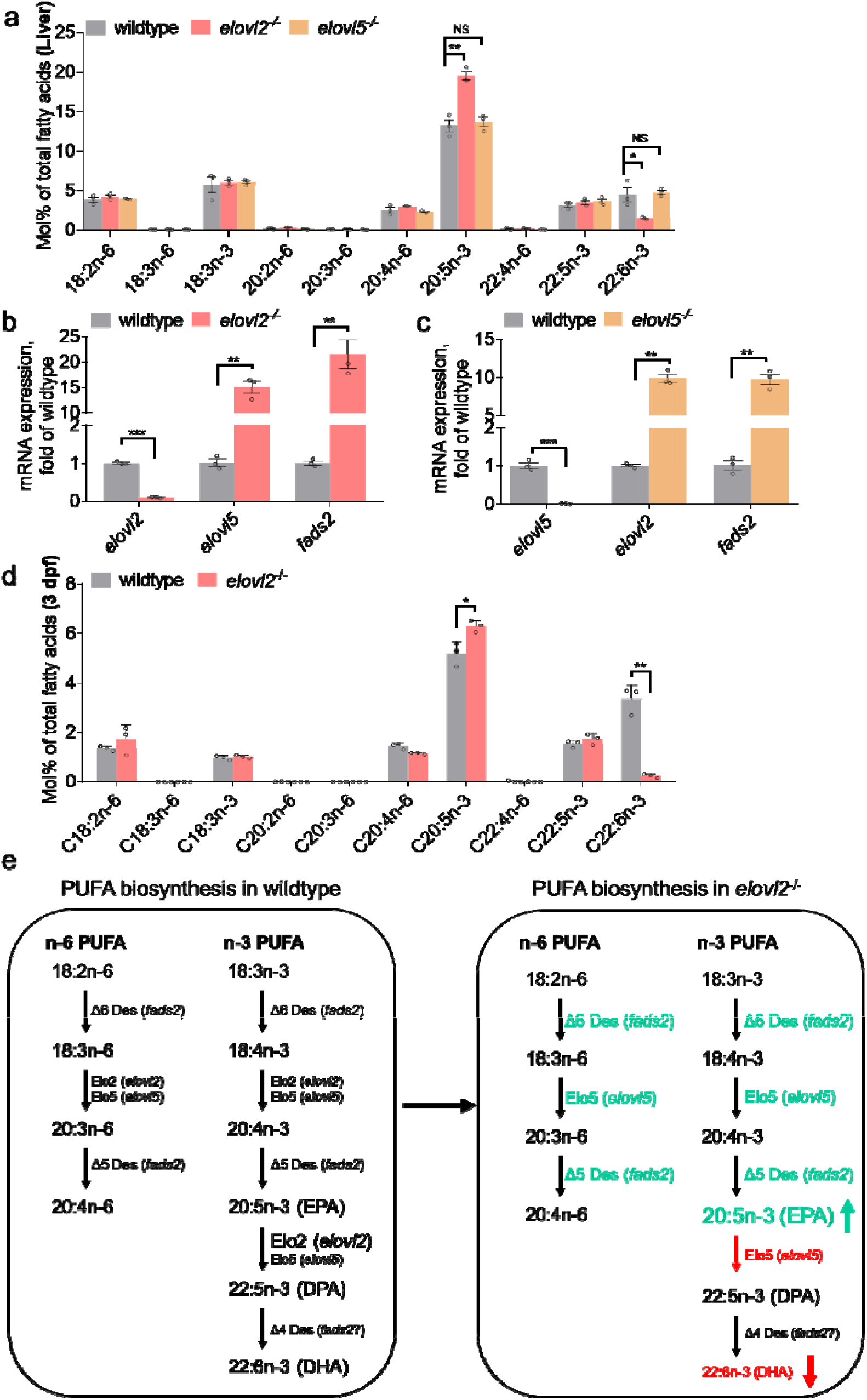
Elovl2 but not Elovl5 is the major elongase mediating the biosynthesis of DHA from EPA in zebrafish. (a) Fatty acid composition (molecular percentage) in the livers of WT (13.15±0.77% for 20:5n-3; 4.42±0.93% for 22:6n-3), *elovl2*^*−/−*^ (19.49±0.57% for 20:5n-3; 1.45±0.09% for 22:6n-3) and *elov5*^*−/−*^ (13.67±0.61% for 20:5n-3; 4.70±0.28% for 22:6n-3). n = 3 replicates. (b) Relative mRNA levels of *elovl2*, *elovl5* and *fads2* in the livers of WT and *elovl2*^*−/−*^ fish examined by RT-qPCR. n = 3 replicates. (c) Relative mRNA levels of *elovl5*, *elovl2* and *fads2* in the livers of WT and *elovl5*^*−/−*^ fish examined by RT-qPCR. n = 3 replicates. (d) Fatty acid composition (molecular percentage) in the embryos of WT (5.18±0.28% for 20:5n-3; 3.34±0.33% for 22:6n-3) and *elovl2*^*−/−*^ (6.29±0.14% for 20:5n-3; 0.23±0.05% for 22:6n-3) at 3 dpf. n = 3 replicates. (e) Diagram showing the biosynthetic pathway of n-6 and n-3 LC-PUFAs in WT and *elovl2*^*−/−*^ zebrafish. The green and red texts indicate the enzyme activities are stimulated or blocked in the *elovl2*^*−/−*^ zebrafish. The green and red arrows indicate the increase or decrease of the content of EPA and DHA in *elovl2*^*−/−*^ zebrafish. All values are the mean ± SEM. Student’s t-test was used in all panels. *, *P* < 0.05, **, *P* < 0.01, *** *P* < 0.001., and NS, no significant difference.

To explore the reasons for the different fatty acid composition between two mutants, we compared the expression level of genes in the PUFAs synthesis pathway in the liver of WT and two mutants. Interestingly, in the *elovl2*^*−/−*^ liver, the expression of *elovl2* was nearly absent, while the expression levels of *elovl5* and *fads2* were significantly increased by 15.1-fold and 21.5-fold, respectively (Fig. 2b). Similarly, in the liver of *elovl5*^*−/−*^ adult, expression of *elovl5* was nearly absent, while the expression levels of *elovl2* and *fads2* were both increased by 9.9-fod and 9.7-fold, respectively (Fig. 2c). These results indicate that the transcription of *elovl2* is activated in *elovl5* mutant and vice versa, however *elovl2* is able to fully compensate the genetic loss of *elovl5*, but even high amount of Elovl5 could not substitute the endogenous enzyme activity of Elovl2.

We further confirmed the role of Elovl2 in the endogenous synthesis of DHA in the *elovl2*^*−/−*^ embryos at 3 dpf, in which the content of DHA was significantly decreased by 93%, and the content of EPA was significantly increased by 21% (Fig. 2d). This further validate the endogenous function of *elovl2* in the embryonic developmental stage and the *elovl2*^*−/−*^ zebrafish is an ideal DHA-deficient model.

## Discussion

Previous studies have shown that both fish Elovl2 and Elovl5 display elongation activities toward C18 and C20 PUFAs in in vitro yeast system (Lebold et al., 2011, Monroig et al., 2009, Agaba et al., 2004). In our study, we have utilized CRISPR/Cas9 technology to knock out both *elovl2* and *elovl5* in zebrafish, therefore clearly clarify the distinct endogenous elongase activity of Elovl2 and Elovl5 by using those in vivo genetic models. We prove that Elovl2 but not Elovl5 is required for endogenous conversion of C20 EPA to C22 docosapentaenoic acid (DPA, 22:5n-3), therefore *elovl2* mutants show deficiency of DHA synthesis.

In vertebrates, the endogenous synthesis of DHA may go through two alternative pathways. In mammals, the endogenous synthesis of DHA is mediated by the Sprecher pathway (Sprecher, 2000), in which EPA is converted to DPA and then 24:5n-3, and 24:5n-3 is subsequently desaturated and subjected to chain shortening by partial β-oxidation, leading to production of DHA. Mammalian Elovl2 is required for elongation of DPA to 24:5n-3, thus depletion of *elovl2* in mammals led to deficiency of DHA and accumulation of DPA (Gregory et al., 2013, Pauter et al., 2014). In teleost fish, however, it is proposed that both the Sprecher pathway and the Δ4 pathway are active for DHA synthesis (Oboh et al., 2017, Li et al., 2010). In the Δ4 pathway, DPA is directly desaturated by Δ4 desaturase to yield DHA. Unlike what was observed in the *elovl2*^*−/−*^ mammals (Gregory et al., 2013, Pauter et al., 2014), we did not detect an accumulation of DPA, the substrate of Sprecher pathway in the *elovl2*^*−/−*^ zebrafish. Instead, the substrate of Δ4 desaturase - EPA is strongly accumulated in the *elovl2*^*−/−*^ zebrafish. Therefore, our study strongly indicates that the Δ4 pathway should be the major pathway for endogenous synthesis of DHA in zebrafish.

Recent studies have established a concept of genetic compensation, in which the genetic disruption of one gene might trigger the upregulation of other genes with sequence similarity, through a machinery of premature termination codon (PTC) mediated nonsense mRNA decay (Ma et al., 2019, El-Brolosy et al., 2019). We noticed that PTCs exist in the coding sequences of elongase domains in both *elovl2* and *elovl5* mutant alleles (Fig. 1c) and both genes share high sequence similarity (data not shown), therefore the strong upregulation of *elovl5* in *elvol2* mutants and vice versa were likely due to the mechanism of genetic compensation in certain mutants. Given that the fatty acid profile in the *elovl5*^*−/−*^ liver was comparable to that in WT liver, we speculated that the upregulation of *elovl2* and *fads2* in the *elovl5*^*−/−*^ liver could completely compensate the genetic depletion of *elovl5*. However, although the *elovl2* mutant showed a dramatic upregulation of *elovl5*, it still displayed a significant deficiency of DHA. This indicates that the ectopically upregulated Elovl5 could not replace the endogenous elongase function of Elovl2 toward the C20 fatty acid, EPA. All these confirm that Elovl2, but not Elovl5, is the main elongase for endogenous synthesis of DHA from its precursor, EPA.

Overall, based on the present study by generating zebrafish mutants of *elovl2* and *elovl5*, both endogenous Elovl2 and Elovl5 present elongase activity toward C18 and C20, but Elovl2 is the major elongase mediating the synthesis of DHA from EPA via the Δ4 pathway in zebrafish (Fig. 2e). In future, the *elovl2*^*−/−*^ zebrafish could be used as an ideal DHA-deficient model to study the endogenous function of DHA.

